# Robust classification of Immune Subtypes in Cancer

**DOI:** 10.1101/2020.01.17.910950

**Authors:** David L Gibbs

## Abstract

As part of the ‘immune landscape of cancer’, six immune subtypes were defined which describe a categorization of tumor-immune states. A number of phenotypic variables were found to associate with immune subtypes, such as nonsilent mutation rates, regulation of immunomodulator genes, and cytokine network structures. An ensemble classifier based on XGBoost is introduced with the goal of classifying tumor samples into one of six immune subtypes. Robust performance was accomplished through feature engineering; quartile-levels, binary gene-pair features, and gene-set-pair features were computed for each sample independently. The classifier is robust to software pipeline and normalization scheme, making it applicable to any expression data format from raw count data to TPMs since the classification is essentially based on simple binary gene-gene level comparisons within a given sample. The classifier is available as an R package or part of the CRI iAtlas portal.

**Code / Tool availability:** Source Code https://github.com/Gibbsdavidl/ImmuneSubtypeClassifier

Web App Tool https://www.cri-iatlas.org/

## Introduction

In the immune landscape paper, six immune subtypes were defined which describe observed immune states found in tumor microenvironments [1]. A number of phenotypic variables were found to associate with immune subtypes, such as nonsilent mutation rates, regulation of immunomodulator genes, and cytokine network structures. These subtypes were derived from fitting a clustering model on gene set scores (signatures) computed using TCGA Pan-Cancer Atlas gene expression.

An early goal following the immune landscape paper was to enable researchers to classify any submitted gene expression data. However, even with publically available scripts that recapitulated the published results, other sources of data were leading to puzzling results that were clearly wrong.

The problem was in the methods used to compute gene signatures. Some signatures were simply averages over a set of genes for each sample. If a new data set contained expression values that were comparatively lower than TCGA Pan-Cancer expression values, the previously fit clustering model would misinterpret the inputs. Considering the many permutations of software pipelines, preprocessing, and normalization possible, the precomputed clustering model was no longer applicable. Demonstrating the inputs possible, gene expression values for CXCR4, computed using two different software pipelines on TCGA samples, are shown in Figure 1.

**Figure 1.**
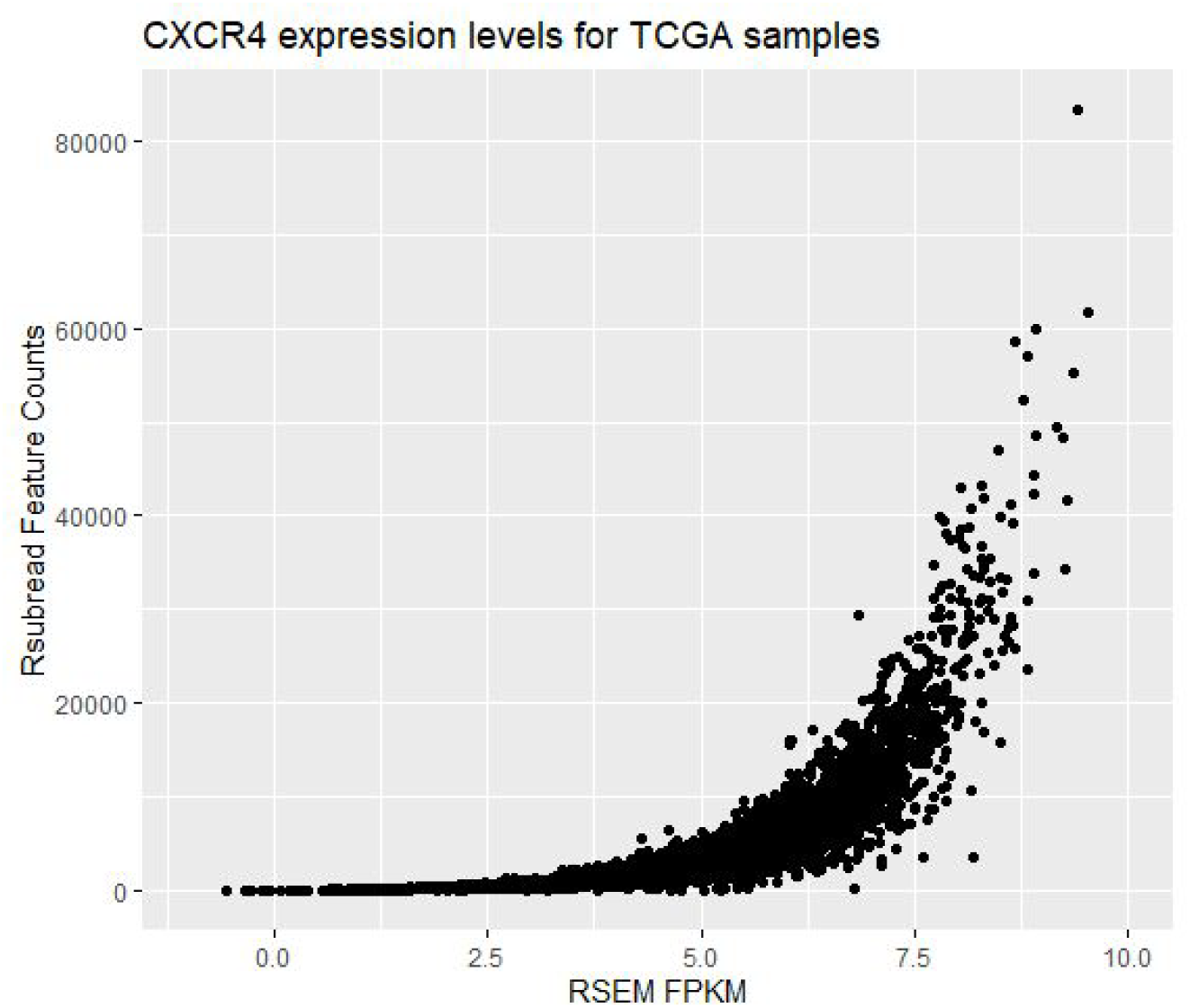
Gene expression values for CXCR4 quantified using Rsubread as feature counts (y-axis) compared to gene quantified using RSEM normalized as FPKM (x-axis) where each point represents one of 7,419 TCGA samples.

Using TCGA data processed in alternative software pipelines, experiments were done to attempt forcing expression distributions to match the TCGA Pan-Cancer data, but ultimately, it was unsuccessful. The classified samples compared poorly to those reported in the landscape manuscript (See Appendix 1).

Instead, a robust classification solution was created that did not depend on data distributions, statistical centers or scales. This solution was based on the prior work of top scoring pairs classifiers [2–5].[6]

This type of classifier has previously been used in cancer research, classifying samples into molecular subtypes which has clear therapeutic advantages [7,8]. Modifications in the models and algorithms have led to improved performance, these include using ensembles of classifiers, decision trees, and feature selection using powerful techniques like Bayesian statistics [9,10].

In ‘top scoring pairs’, a gene-pair (*g*_*1*_ and *g*_*2*_) feature is generated per sample where the feature will take the value *1* if *g*_*1*_ > *g*_*2*_ and *0* otherwise. Attempting classification with all possible gene-pair features is inefficient, making some preliminary feature selection necessary. In this work, genes were selected using the complete set of genes that were used in the derivation of immune subtypes (485 genes).

However, since the original immune subtype work was based on the scores of gene sets, an additional feature was engineered based on gene-set-pairs. This feature takes a value between 0 and 1 which indicates the proportion of *g*_*i*_ > *g*_*j*_ relationships observed where *g*_*i*_ ∈ gene-set_1_ and *g*_*j*_ ∊ gene-set_2_. Additionally, single gene features were created by binning genes, within samples, into quartiles, giving the feature a categorical value.

In summary, for each sample independently, we have features that are gene quartiles (*g*_*1*_-75%), gene-pairs (value of 1 since *g*_*1*_ > *g*_*2*_), and gene-set-pairs (0.765 for *set_1_ > set_2_*). Each sample is associated with a vector of feature values.

## Results

We used the XGBoost classifier [11], with binary gene-pair, fractional gene-set-pair, and quantile features, to classify each subtype independently. Batch corrected gene expression data from the TCGA Pan-Cancer Atlas with 9,129 samples were used to train the model [1,12]. First six binary classifiers were trained, predicting inclusion / exclusion of each subtype (In-C2 vs. Not-In-C2). The resulting scores from the six classifiers were then used to train one additional XGBoost classifier to make the ‘final call’. XGBoost parameters are found in Appendix 2. A gene filtering step based on mean rank differences between groups was used, and cross validation was used to determine the number of XGBoost trees.

To reduce variance in the classification, an ensemble approach was applied. Each member of the ensemble was created by randomly sampling 80% of the available samples. The ensemble consisted of 10 classifiers for each subtype, each of which return a probability. The probabilities for each subtype classifier (C1-vs-not-C1) are averaged across the ensemble, resulting in six probabilities (C1-C6) for each input sample. This table, of one call per binary-subtype-classifier, is fed into a final XGBoost classifier that was trained on immune subtypes, to produce a final ‘best call’ output. See Figure 2.

**Figure 2.**
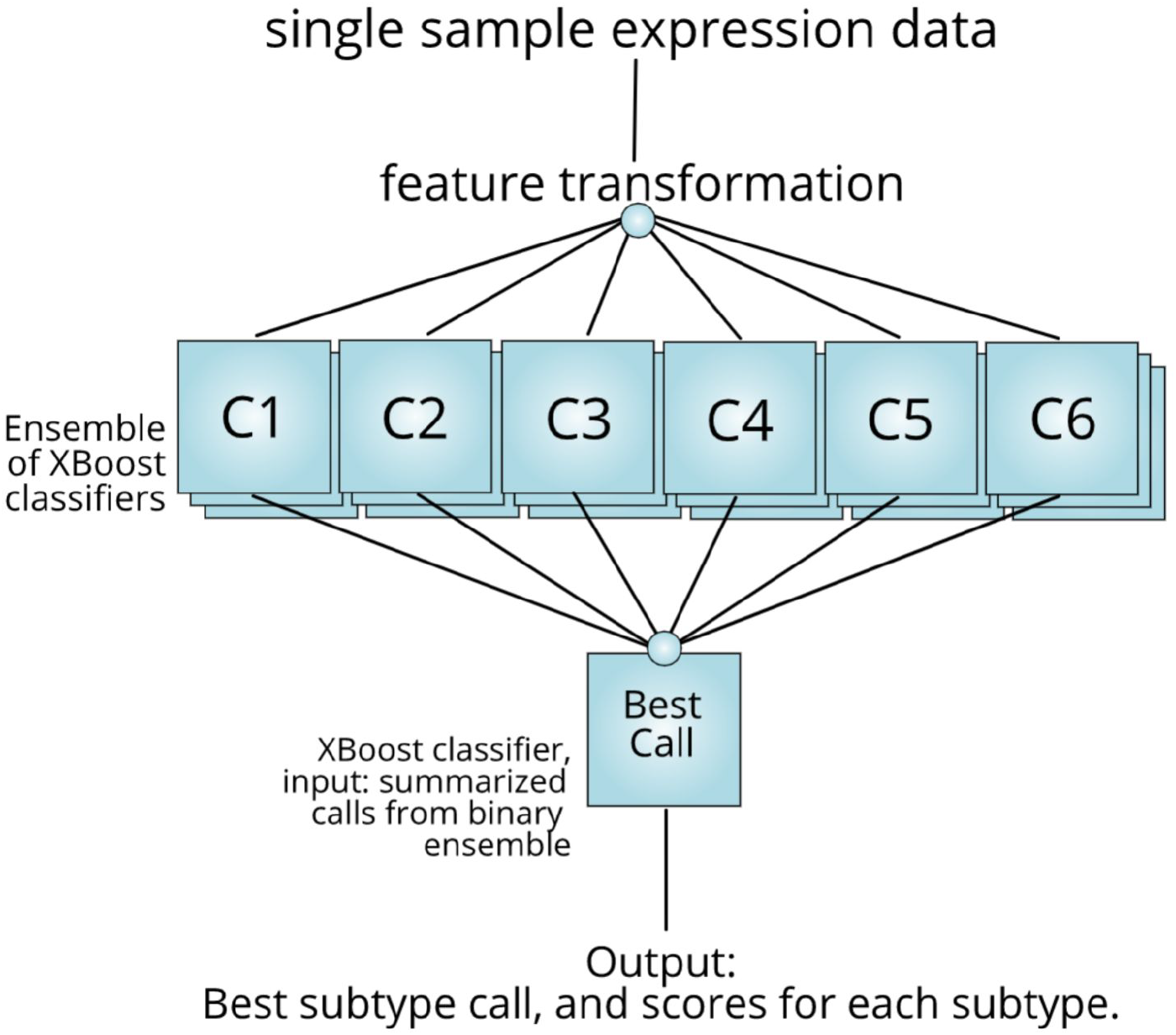
The classifier is composed of a number of parts, each trained with immune subtypes from the ‘Immune Landscape of Cancer’ manuscript. For each sample, features are computed which include single gene quartile levels, binary valued gene-pair features, and numeric gene-set-pair features. The computed features are input to an ensemble of XGBoost classifiers, each making a prediction on a single subtype. The outputs are averaged and used as input to a final classifier used to make a ‘best call’.

Five data sets, representing different processing pipelines, were used to test the classifier performance. Each pipeline had been run on barcoded TCGA samples, producing matched sample data. Software includes Kallisto (transcript-level TPM), RSEM (gene level FPKM & TPM), and Rsubread (gene level feature counts) [13–16]. Data was downloaded from XENA Hub and NCBI GEO (GSE62944) [17,18]. The Kallisto transcript level values were summed to gene level values, using the gencode v23 transcript map [19].

In all cases, genes were not normalized across samples, for example by median centering. Since the classifier is based on gene-gene relationships within a sample, by performing normalization across samples, the relationship can be highly affected leading to altered results.

<u>Therefore, it is absolutely vital not to attempt any such normalization that makes use of information across samples.</u>

Classification results are shown in Table 1. For each subtype, the precision (TP / TP+FP) and the recall (TP / TP + FN) are listed. Also, the total accuracy across all subtypes is listed (diag TP / total samples). All data sets show accuracies above 90%.

Using XGBoost, it is possible to rank the information gain of features, which in this case, are quartiles, gene-pairs, and gene-set-pairs. Information gain was summed, within subtype classifier and across ensembles, to get a single table of features per subtype. A full table is available in the github repository. Overall, the gene-set-pair features were most informative, followed by gene-pairs, with a few single-gene-quantiles also in there. Top features for each subtype are shown in Table 2. Signature-features are found in Appendix 3.

| Software | Norm | Samples | Precision / Recall |  |  |  |  |  | Total acc. |
| --- | --- | --- | --- | --- | --- | --- | --- | --- | --- |
|  |  |  | C1 | C2 | C3 | C4 | C5 | C6 |  |
| RSEM | FPKM | 8500 | 98.3<br>97.7 | 97.1<br>99.9 | 98.1<br>99.3 | 99.6<br>91.9 | 99.7<br>99.2 | 92.4<br>94.4 | 98.0 |
| RSEM | TPM | 8500 | 98.3<br>97.6 | 97.1<br>99.8 | 97.9<br>99.3 | 99.6<br>91.9 | 99.7<br>99.2 | 92.4<br>94.4 | 98.0 |
| Kallisto<br>(transcripts) | TPM | 8500 | 97.0<br>98.3 | 99.4<br>98.4 | 98.1<br>98.0 | 96.5<br>97.5 | 98.2<br>98.9 | 93.3<br>85.0 | 97.9 |
| Rsubread | TPM | 6680 | 91.4<br>90.8 | 87.9<br>98.1 | 92.0<br>93.2 | 97.1<br>73.7 | 98.0<br>93.7 | 69.9<br>59.6 | 91.1 |
| Rsubread | Counts | 8037 | 91.7<br>93.8 | 94.5<br>96.2 | 92.6<br>94.6 | 97.1<br>82.3 | 97.3<br>98.1 | 70.3<br>73.1 | 93.2 |

**Table 2.** Top 3 predictive features for each subtype. The sum over the ensemble members gives the total information gain of a given feature. Features in the form of ‘sNsM’ represent a feature computed from the pairwise comparison of genes in signature N and signature M where signatures are those used to derive the immune subtypes. A pair of gene, separated by a colon, are a binary gene-pair feature, and a single gene name represents a quartile feature.

| Subtype | Feature | Gain_sum |
| --- | --- | --- |
| 1 | s3s4 | 2.716080974 |
| 1 | MX1:MYBL2 | 2.304420705 |
| 1 | s2s4 | 1.224188047 |
| 2 | s3s4 | 1.79241718 |
| 2 | GNG11:OASL | 2.042958694 |
| 2 | OAS2:RHOQ | 1.787712903 |
| 3 | s1s2 | 1.255178685 |
| 3 | s2s3 | 1.131509752 |
| 3 | MYBL2 | 3.7697349 |
| 4 | s3s4 | 1.143575419 |
| 4 | s1s2 | 1.512503626 |
| 4 | CPEB4:COL6A3 | 1.063994637 |
| 5 | TIMP1:RASSF2 | 5.863580618 |
| 5 | TIMP1:EVI2A | 2.391847181 |
| 5 | COL3A1:SELL | 1.085996179 |
| 6 | s3s4 | 0.8408236112 |
| 6 | s2s5 | 1.989936177 |
| 6 | s4s5 | 1.172035395 |

## Discussion

The initial work used to create immune subtypes, was based on batch corrected TCGA Pan-Cancer gene expression data, which was created with the intention of allowing comparisons across tissue types.

In the ‘Immune Landscape of Cancer’ manuscript, gene sets were scored on a single sample basis, essentially performing dimension reduction, where 20,000+ genes were replaced with functional scores. The scoring methods were taken from a collection of literature and varied considerably, where one method might be simply taking the mean, while another used precomputed *eigengenes* from a WGCNA type analysis [20]. Computing the gene set scores created a new matrix (tumors by gene set scores) that was used as an input to clustering.

However, the commonly used method of consensus clustering was infeasible with 10K samples due to time and memory constraints [21]. Therefore, mclust, a model based clustering method was used. Once the model is fit, it can be used to make predictions on new samples [22,23].

Mclust models represent a mixture of gaussians and with a very long matrix with five gene sets in columns and 9K rows, it presents an ideal situation for fitting a statistical model. We concluded with a small study showing that pulled out samples did not change the model and could be predicted, confirming some robustness in the clusters.

However, it was discovered that the model was highly sensitive to the method of gene quantification. In order to call immune subtypes on new samples, a new method was required. Thus, we started working towards a scheme that would be completely independent of software pipeline, independent of expression magnitudes. This led us to looking at robust feature engineering using quantiles, gene-gene relationships, and eventually set-set relationships. The computed features were uncoupled from the magnitude of expression, and were found to be very capable of distinguishing immune subtypes.

## Conclusion

A classification model has been introduced that is robust to the variation introduced by different software and statistical pipelines by engineering features derived from ‘top scoring pairs’ style classification. Additionally, beyond gene pairs, we also introduce gene-set-pairs and gene-quartile features that are found in the lists of most informative features. The tool is available as an R package and part of the CRI-iAtlas web app.

## Acknowledgements

Funding from the Cancer Research Institute is gratefully acknowledged. Special thanks to the iAtlas working group and Vincent Lavergne at the Unicancer research center in Lyon, France.

## Data Used

### Training data

https://gdc.cancer.gov/about-data/publications/pancanatlas (RNA (Final) - EBPlusPlusAdjustPANCAN_IlluminaHiSeq_RNASeqV2.geneExp.tsv)

### Testing data

UCSC Xena hub ID: tcga_RSEM_gene_fpkm, version: 2016-09-01

UCSC Xena hub ID: tcga_RSEM_gene_tpm, version: 2016-09-01

UCSC Xena hub ID: tcga_Kallisto_tpm, version: 2019-02-25

NCBI GEO: GSM1536837 1-27-15 TPM

NCBI GEO: GSM1536837 06-01-15 FeatureCounts

## Appendix 1.

Results on a subset of 1000 samples of RSEM FPKM data using the ‘Immune-Landscape’ mclust model. New cluster labels were manually aligned, but shows poor performance. mclust predictions are on the columns (calls) and true calls (from immune landscape manuscript) are on rows.

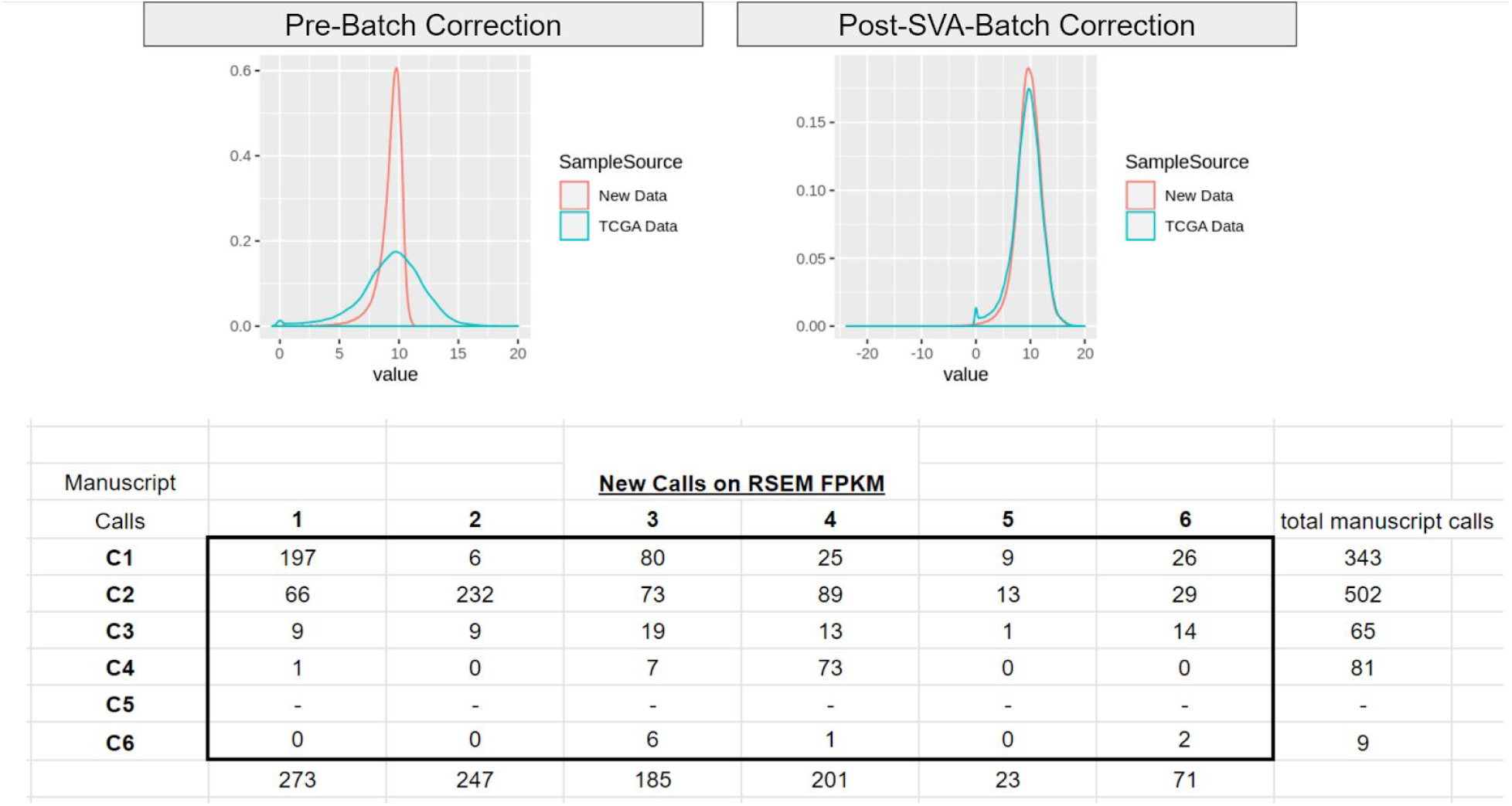

## Appendix 2. XGboost classifier fitting parameters

max_depth = 5 # decision tree depth

eta = 0.3 # learning rate

nrounds = 150 # total rounds used in CV

nfold=5 # fold size in CV for selecting XGboost rounds

ensemble_size=10 # number of ensembles

sampSize=0.8 # ensemble sample size

breakVec=c(0, 0.25, 0.5, 0.75, 1.0) # quartile levels

ptail=0.01 # gene filtering

## Appendix 3. Getting the names of the signatures

~~~
library(ImmuneSubtypeClassifier)
data(“geneSetSymbols”)
names(genesetsymbols)
1. “LIexpression_score”
2. “CSF1_response”
3. “Module3_IFN_score”
4. “TGFB_score_21050467”
5. “CHANG_CORE_SERUM_RESPONSE_UP”
~~~

## Getting the names of genes used in training the classifier

~~~
data(“modelgenes”)
modelgenes
~~~

## Getting the full feature information-gain table

https://github.com/Gibbsdavidl/ImmuneSubtypeClassifier/blob/master/inst/important_features_in_the_ensemble_model.tsv

